# Chromonomer: a tool set for repairing and enhancing assembled genomes through integration of genetic maps and conserved synteny

**DOI:** 10.1101/2020.02.04.934711

**Authors:** Julian Catchen, Angel Amores, Susan Bassham

## Abstract

The pace of the sequencing and computational assembly of novel reference genomes is accelerating. Though DNA sequencing technologies and assembly software tools continue to improve, biological features of genomes such as repetitive sequence as well as molecular artifacts that often accompany sequencing library preparation can lead to fragmented or chimeric assemblies. If left uncorrected, defects like these trammel progress on understanding genome structure and function, or worse, positively mislead such research. Fortunately, integration of additional, independent streams of information, such as a genetic map – particularly a marker-dense map from RADseq, for example – and conserved orthologous gene order from related taxa can be used to scaffold together unlinked, disordered fragments and to restructure a reference genome where it is incorrectly joined. We present a tool set for automating these processes, one that additionally tracks any changes to the assembly and to the genetic map, and which allows the user to scrutinize these changes with the help of web-based, graphical visualizations. Chromonomer takes a user-defined reference genome, a map of genetic markers, and, optionally, conserved synteny information to construct an improved reference genome of chromosome models: a “chromonome”. We demonstrate Chromonomer’s performance on genome assemblies and genetic maps that have disparate characteristics and levels of quality.

## Introduction

Researchers are generating new reference genomes at an accelerating pace. While it is now trivial to produce enough sequence information to cover even large genomes many times over, the assembly of a realistic reference genome can still be challenging for both bioinformatic and biological reasons (Ghurye and Pop 2019; Church et al. 2011; Nowoshilow et al. 2018; De La Torre et al. 2014). A high-quality reference genome with minimized gaps and misassemblies, particularly one organized into chromosomes – known as a *chromonome* (Braasch et al. 2015) – is a valuable research tool. Comparative genomics studies that have employed, for example, the analysis of conserved synteny of genes among distantly-related taxonomic groups has led to better understanding of how genes and genomes evolve and function (Naruse 2004; Jaillon et al. 2004; Shah et al. 2012; Lovell et al. 2014; Zhao and Schranz 2019). Likewise, to understand the population dynamics of selection and drift, as described by measures of mutation and linkage, requires chromosome-level stretches of sequence (Hohenlohe et al. 2010; Luikart et al. 2018). Reliably assembled reference genomes have aided exploration of chromosome structural conservation or rearrangement through evolutionary time (Wang et al. 2013; Jay et al. 2018), the effects of transposable element perturbation (Woronik et al. 2019), the fate of duplicated genes following divergence of organismal lineages (Brunet et al. 2006; Kassahn et al. 2009), the mechanisms of long distance regulation of genes (Kleinjan and van Heyningen 2005), and the progression of disease-resistant alleles in populations (Epstein et al. 2016). New reference genomes that are misassembled, or that remain broken in scaffolds, or whose scaffold order relies on the reference genome of a different taxon, can stall inferences about critical biological processes or, worse, can mislead.

Construction of a genetic map is still relevant for a variety of research goals; for example, comparing a genetic map with a physical genome sequence aids identification of gene candidates causal for variant or mutant phenotypes (Peichel and Marques 2017; Meinke et al. 2003), and reveals variation in recombination rate across the genome (Roesti et al. 2013; Dukić et al. 2016), an important consideration in population genomics or genome-wide association studies. In addition, a map can also benefit the assembly of a reference genome in several valuable ways. A map can help reveal points of artifactual contiguity in an assembly, can bind scaffolds into “linkage groups” that are chromosome models, and can order and orient the scaffolds relative to one another. It is now relatively straightforward and rapid to genotype individuals at thousands of loci by using one of many massively parallel sequencing methods such as Restriction site-associated DNA sequencing (RADseq; (Baird et al. 2008; Andrews et al. 2016; Davey et al. 2011)). Marker-dense maps have the potential to capture a majority of the assembled genome length into linkage groups. Perhaps more importantly, a genetic map provides an independent line of evidence to verify the genome assembly itself. Potentially chimeric scaffolds can be detected where the physical and genetic map relationships of markers on scaffolds conflict, such as in cases where a single scaffold’s markers map to more than one linkage group. The efficacy of a genetic map for consolidating an assembly and validating its quality depends on a number of important factors, including the density and distribution of markers, the number of meiotic crossovers represented in the mapping cross progeny, the size distribution of the scaffolds, and the granularity of misassembly with respect to the distance between markers.

A large number of scaffolds often results from assembly of complex genomes, particularly when short-read genome sequencing data are used. Assemblies based even on long-read technology that provide robust assembly lengths can still contain major assembly errors stemming from the presence of complex repeats or through the application of software that hybridizes multiple assemblies together (Sohn and Nam 2016). Making hand corrections to a genome assembly via a comparison of genetic and physical maps is prohibitively time-consuming. In the end, a genome assembly is a hypothesis that proposes a sequence order while the true order will always remain unknown. A useful tool should be able to automate the flagging of problematic scaffolds, resolve conflicts between the assembly and the genetic map in a rational and efficient way, and integrate additional lines of evidence that support a hypothesis of genomic structure. We present here software we call Chromonomer that corrects, orders, and orients scaffolds by integrating genetic maps and genome assemblies. Chromonomer can create chromosome-level assemblies while providing extensive documentation of how the elements of evidence fit together. To further improve assemblies, the software can integrate auxiliary lines of evidence such as conserved gene synteny and raw read depth of coverage, and it provides tools to extract gene annotations from a scaffold-level assembly and translate their locations to a chromosome-level assembly (and vice versa). Earlier, prototype versions of Chromonomer have been used in a number of published genome assembly integrations (e.g., Amores et al. 2014; Fountain et al. 2016; Small et al. 2016; Kim et al. 2019; Moran et al. 2019). Here we illustrate the performance of Chromonomer with three qualitatively different test cases in the teleost reference genomes: a high-quality, short-read-based assembly with a map made from a modestly sized genetic cross (Gulf pipefish), a high quality, long-read-based assembly with a large genetic cross (platyfish), and a highly scrambled, short-read-based assembly with a large genetic cross (Antarctic black rockcod).

## Results

As sequencing technologies have evolved, the types of information they provide about the underlying DNA sequence and their respective error models have changed, resulting in shifting assembly strategies. Genome assembly has progressed through three major strategies since the inception of high-throughput sequencing: short-read-only assemblies, hybrid assemblies that incorporated longer reads to join and gap-fill short-read assemblies, and long-read-only assemblies.

The first massively parallel sequencers, such as the Illumina HiSeq machines (Reuter et al. 2015), were short-read technologies. Two major molecular library types most useful for genome assembly could be sequenced with this technology: first, the two ends of sheared genomic DNA molecules could be sequenced (paired-end shotgun libraries) from fragments 200bp to 1Kbp long. Second, much longer molecules, from 2.5Kbp to 20Kbp could be circularized and sheared in the generation of “mate-pair” libraries, with the goal of sequencing the circularized junction to connect distant sequences. Assemblers designed around this technology used a *De Bruijn* graph structure to build gapless *contigs* from k-mers extracted from the reads (Compeau et al. 2011). Next, mate-pair reads would be overlain on the contigs, and the contigs could be ordered and oriented into *scaffolds*, separated by gapped sequence that was unknown but inferred from the mate-pair reads using the approximate original DNA fragment length (Gnerre et al. 2011; Luo et al. 2012; Chapman et al. 2011). Since contigs are derived from connected k-mers that were sequenced from unmanipulated molecules, they are generally reliable. Mate-pair libraries can have a much higher error rate, however, as the circularization and shearing possibly creates chimeric reads, which can result in scaffolds that are incorrectly joined. Mate-pair reads that happen to land in repeats of different lengths can generate additional assembly error in order and orientation.

Initially, long-read sequencing was expensive and low volume, but could be employed to augment the scaffolds of short-read assemblies. Read lengths from 400bp (454 GLX II sequencing; Pop, (2009)) up to 10-15Kbp (PacBio RSII and Oxford Nanopore; Reuter, et al. (2015)) could be integrated into short-read assemblies with software such as PBJelly (English et al. 2012), QuickMerge (Chakraborty et al. 2016), and many others. While these hybrid assemblies have improved quality metrics like N50 and L50, they add a second layer of scaffolding on top of the short-read assembly, subject to similar issues when sequence repeats create artifactual joining by partial sequence matches.

Long-read sequencing technologies, particularly with the introduction of the Sequel I and II platforms (Pacific Biosciences), now have high enough volume that new assemblies are being constructed purely from long reads. Although long reads currently have high indel-derived error rates, those errors can be corrected with sufficient sequencing depth (Fu et al. 2019). These assemblies have huge N50 measures, and typically contain very few or even no gaps because very limited or no scaffolding is employed, although repeats occasionally result in mis-joins of contigs.

To further increase assembly contiguity, approaching chromosome-level, an additional set of technologies has been employed, including optical maps (Howe and Wood 2015; Pendleton et al. 2015) and Hi-C libraries (Lieberman-Aiden et al. 2009). Optical mapping relies on restriction enzymes that nick and label high molecular weight DNA followed by high-resolution detection of the pattern of cut sites. There is a degree of error in translating physical cut site patterns back to the sequenced location of cut sites in the assembled genome. Hi-C libraries are constructed from genomic DNA that is first crosslinked by a fixative while in the nucleus, and then sequenced on short-read platforms. Reads derived from cross-linked DNA fragments can be used to infer genomic segments that are located at a distance but likely on the same linear DNA molecule and are used as a basis to scaffold contigs. As with other scaffolding technologies, these processes can create incorrect joins.

Some of the earliest developed genomic resources include genetic maps, which were used to map the chromosomal locations of some of the first inversion phenotypes and mutants to be described in *Drosophila*, using polytene chromosomes (Painter 1933; Schaeffer et al. 2008). Genetic maps rely on the discovery of markers (visible chromosome bands in the foregoing example) and the tracking of markers as they change haplotypic neighbors through recombination during one or more generations. Genetic mapping has experienced many transitions in marker technology, with the most recent typically based on RAD markers. These SNP or haplotype markers are ordered statistically, relying on the number of observed recombination events between alleles in the cross to derive a measure of distance between them. The end result is a set of linkage groups, where each group represents a chromosome segment or, ideally, a complete chromosome, described by a set of markers ordered in centiMorgan (cM) units of genetic distance. If the markers can be located within a draft genome assembly (as RADseq markers can), genetic maps can provide an independent source of information to define the order and orientation of assembled genomic fragments. While genetic maps can provide very reliable information, precise resolution of marker order is determined by the number of recombination events (in that more events yield better ordering), as well as on genotyping accuracy. Genotyping errors can place markers in the wrong positions on the resulting linkage groups.

The primary design goal of Chromonomer is to integrate assembled contigs and scaffolds with genetic maps to produce a chromosome-level assembly. Each of the above described assembly techniques provides information of varying reliability. For example, mate-pair reads can create a chimeric scaffold during assembly or miscalled genotypes can alter marker positions in a map. Chromonomer seeks to integrate disparate information, using the most reliable information first. In cases where the first source of information is ambiguous, the software can apply additional sources. Chromonomer is designed first to trust contiguous genome assembly (*contigs*, where scaffolding has not yet been inferred from other molecular information). Next, Chromonomer trusts the overall linkage map ordering, followed by the scaffolding, raw read depth of coverage, and finally, conserved gene synteny, depending on what information is available and on the user’s dictate.

Chromonomer requires several components in order to work. First, it needs a description of a physical genome assembly (an AGP – *A Golden Path* – file; (NCBI 2019)). The AGP file is expected to be defined at the scaffold level, with each scaffold described in the file as a set of oriented contigs and gaps. In addition, Chromonomer needs a description of the genetic linkage map (in a tab-separated values file) containing, for each linkage group, the set of markers identified by unique IDs and their respective cM positions. A BAM file (SAM/BAM Format Specification Working Group 2019) supplies the alignment position of each marker within the physical assembly; the alignment information includes the physical sequence of each marker, each labeled with the same ID contained in the genetic linkage map file.

By definition, a *scaffold* is a collection of oriented *contigs* – which themselves are runs of gapless sequence. Long-read assemblers might produce only contigs, but we will use the terms interchangeably here, except when specifically referring to the presence of gaps. Chromonomer first collects all the markers aligned to each independent scaffold (Fig. 1A-C) and identifies scaffolds that are mapped to two or more linkage groups. Since linkage group assignment is statistically very robust, the linkage map is trusted over the physical scaffolding and such scaffolds are split (Fig. 1B) at the nearest scaffold gap (sequence of N characters) between the sets of markers. If multiple gaps exist there, the largest gap is chosen. If there are not enough markers to support a split (user definable, default 2), the conflicting markers are logged and discarded (Fig. 1C).

**Figure 1.**
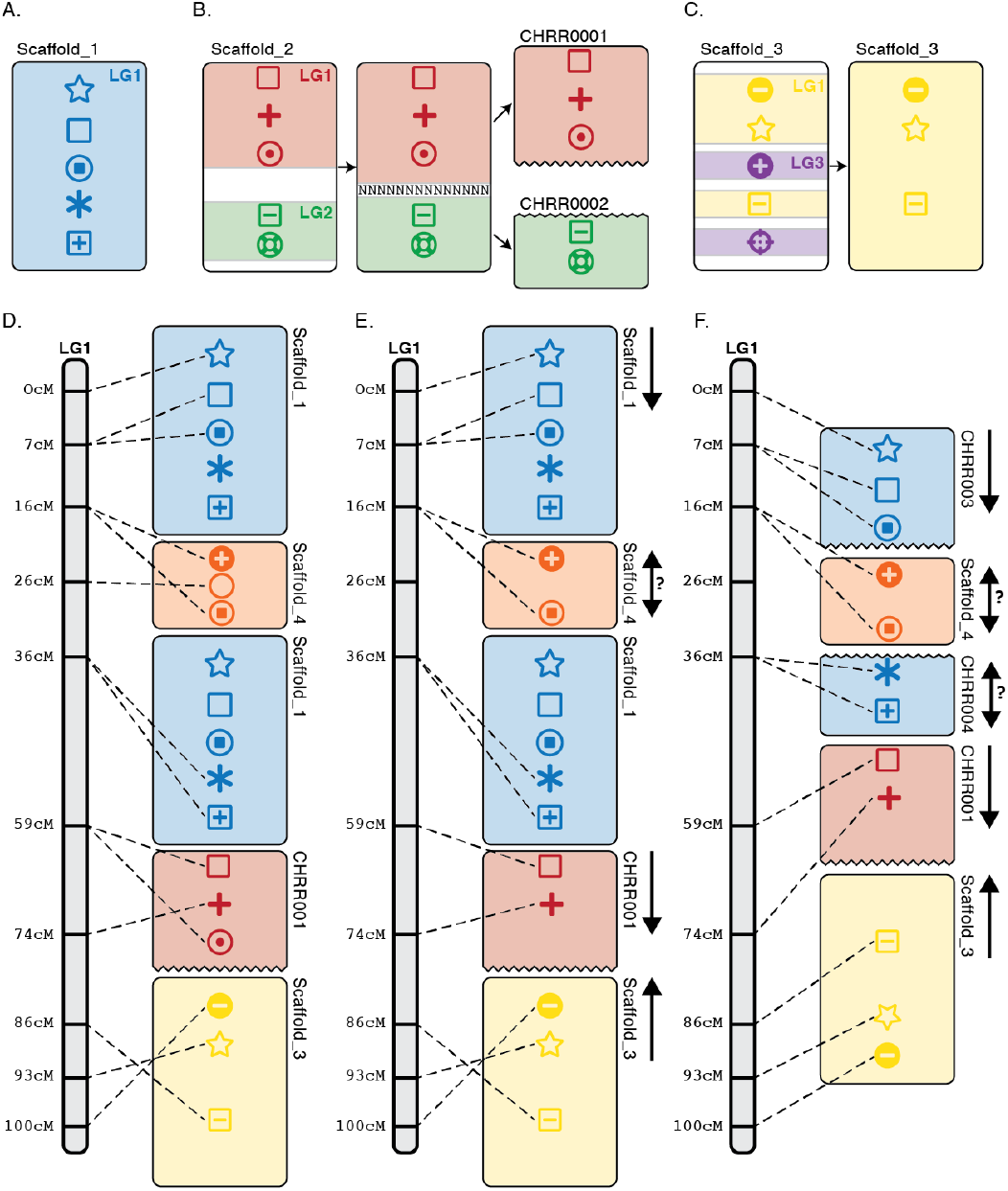
The primary Chromonomer algorithm. The algorithm takes a set of scaffolds and a set of markers, an assembly file (AGP file), which describes how contigs and gaps are formed into scaffolds in the assembly, and a genetic map, which provides the marker order. (A-C) Scaffolds (rounded rectangles) are first evaluated to identify sets of markers (symbols within rectangles) mapped to different linkage groups. Those scaffolds will be split at the nearest gap (B) or pruned out (C) if a consistent set of markers cannot be found. (D) Scaffolds are anchored to their positions in the genetic map, and if a scaffold contains markers from two locations in the genetic map, it is anchored twice. (E) A consistent order of markers is determined, with inconsistent markers discarded. (F) Scaffolds are oriented or split at the nearest gap, as dictated by the genetic map.

Next, Chromonomer constructs a graph to represent each linkage group, in which each cM measure from the linkage group is a node in the graph and each cM node contains one or more markers. Markers are used to anchor scaffolds to linkage groups (using the cM position of each marker and its alignment position in the physical assembly). If a scaffold is anchored to multiple, non-neighboring nodes, it is placed into the graph in every position where at least one of its aligned markers occurs (Fig. 1D). If a scaffold is not anchored to at least two nodes, then it will be impossible from the map alone to determine its “plus” or “minus” orientation relative to the linkage group. If multiple scaffolds collapse into the same, single node, their order (linear series) within the node cannot be determined from the map alone either, though this cluster of scaffolds can still be ordered relative to scaffolds anchored to other nodes. From this it is clear to see how the power to detect misassembly and to order and orient scaffolds depends upon the number of nodes, which is itself a function of the resolution of genetic markers. The more genotyped progeny there are in the cross, the higher the number of potential recombination events that can resolve marker order, and therefore the greater number of nodes and fewer markers per node. Researchers should therefore boost progeny number when feasible.

Unlike how Chromonomer prioritizes map structure over scaffolding, the algorithm trusts the contiguous physical assembly over individual markers that are not corroborated by other, nearby markers. Since genotyping errors can slightly change the position of a particular marker in the map, Chromonomer will search for a sequential set of markers whose physical alignment order is consistent with the map. Out of order markers are discarded. As a scaffold is inserted into the graph at each node where it genetically appears, it is common for a single scaffold to span multiple nodes in the graph and these graph nodes are collapsed. The graph is then searched to find any scaffolds that remain in multiple positions across the graph, a circumstance that indicates that at least one scaffold belongs between these nodes; such duplicated scaffolds are split at their nearest gap boundary (Fig. 1F) to produce the final integration of the map and assembly. Chromonomer logs the markers that were discarded and any scaffolds that were split, listing the conflict and position that caused the split. The software outputs an AGP file describing the integrated assembly and a FASTA file describing the integrated sequence. If requested, Chromonomer can lift over gene annotations to the new assembly coordinates. Chromonomer provides textual outputs as well as a web-based visualization tool; for example, Figure 2 (among others) in this manuscript were modified from outputs initially generated via the web-based visualization.

**Figure 2.**
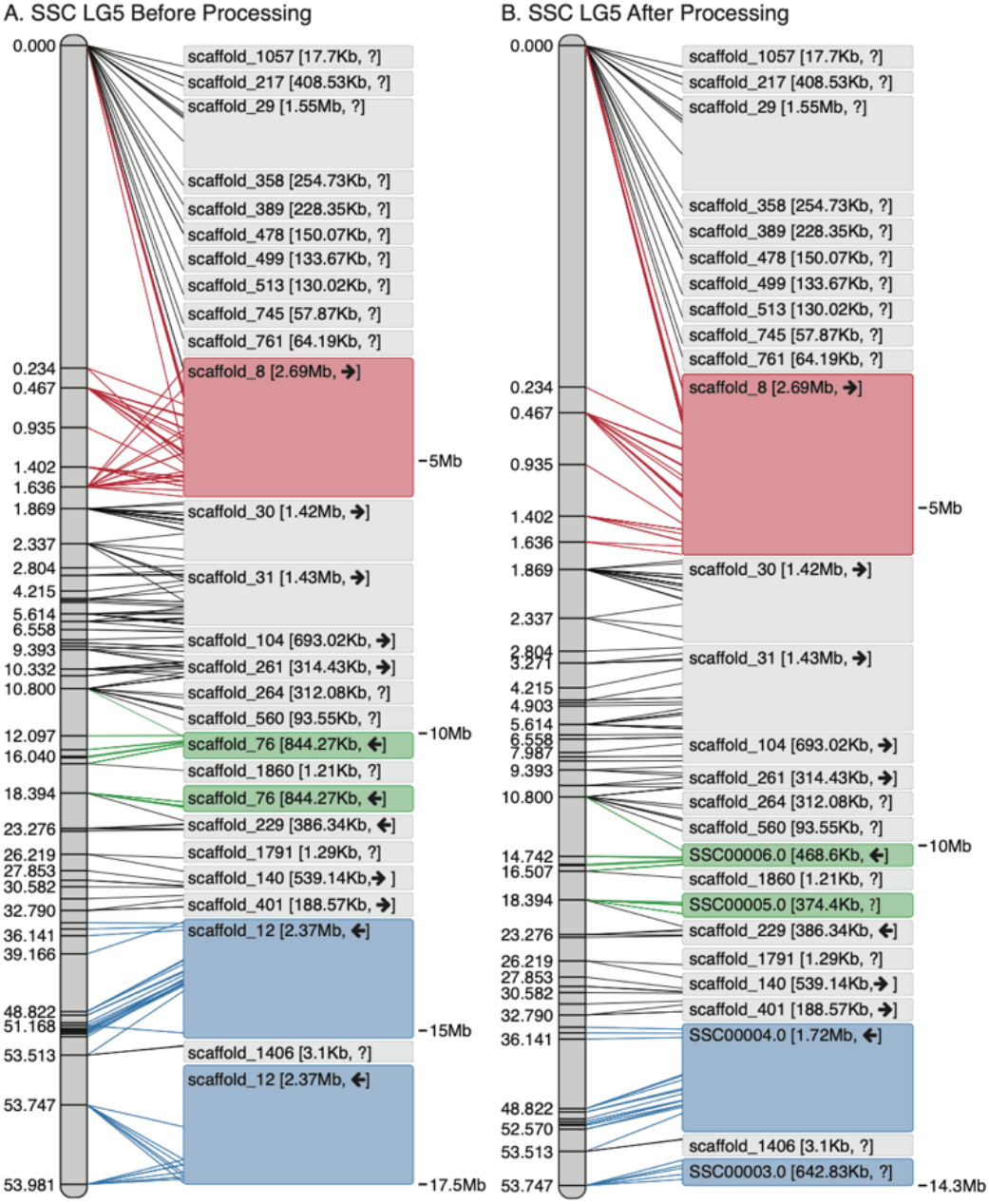
The Chromonomer algorithm as employed in the Gulf pipefish assembly. The figure shows all the numbered scaffolds belonging to LG5 before (A) and after (B) processing. In the diagrams, each marker in the linkage group (left) is connected by a line to its alignment position on each scaffold (right). In red in (A), scaffold_8 demonstrates an example of markers with conflicting physical and map orders. In (B), the order of markers has been resolved and some conflicting markers discarded. Scaffold_76 (green) and scaffold_12 (blue), which are each anchored in two map positions, demonstrate examples of scaffolds that need to be split so a third scaffold can be inserted into the rift.

### The basal Chromonomer algorithm applied to the Gulf pipefish

Chromonomer was used to integrate the physical assembly of the Gulf pipefish (*Sygnathus scovelli*) with a genetic map derived from an F1 cross (Small et al. 2016) of 108 progeny. This reference genome is an Illumina-based assembly following the ALLPATHS-LG assembly strategy (180bp insert length shotgun library with additional mate-pair libraries). The map consisted of 6593 markers on 22 linkage groups while the physical assembly was 307Mbp in total length, contained in 2104 scaffolds, and had a scaffold N50 of 640Kb and an L50 of 115.

Figure 2 shows how the Chromonomer algorithm handled linkage group 5 (LG5). It is clear that prior to processing, marker order is inconsistent with alignment order (e.g., see Fig. 2A, in which red lines are crisscrossed), but after processing their order has been corrected by discarding incongruous markers (the red lines are resolved in Fig. 2B). Scaffold 76 (Fig. 2A, green) appears in the map twice, as does scaffold 12 (Fig. 2A, blue). After processing (Fig. 2B), both scaffolds have been split and, in each case, an additional scaffold has filled a gap (scaffolds 1860 and 1406, respectively). After Chromonomer integrated the genetic map, 266Mbp was incorporated into 22 linkage groups in the chromonomed assembly (87% of assembly length) with 550 of the 2104 scaffolds incorporated and five scaffolds having been split. Of those scaffolds not incorporated, the mean and N50 lengths were 26Kbp and 4Kbp, respectively, and no markers aligned to 1432 of these 1554 relatively small scaffolds.

### Incorporating conserved gene synteny into the Gulf pipefish integration

If a scaffold or contig does not span more than one node in the linkage map, Chromonomer is unable to determine its orientation. Variation in rates of recombination across the genome caused, for example, by the presence of structural variants, sex chromosomes, or having a small number of progeny in the mapping cross can leave a substantial number of scaffolds without a determinable order and orientation. Scaffolds in these regions are oriented as if they were in the forward direction and are ordered arbitrarily within the mapping node to which they are linked. Optionally, Chromonomer can incorporate conserved gene synteny from an external genome to further resolve these scaffolds. The user supplies to Chromonomer 1) a scaffold-level annotation of genes (via a GTF or GFF file) for the focal genome, 2) a chromosome-level annotation of the external, synteny-supplying genome, and 3) a tab-separated file defining orthologous genes between the two genomes. For regions of the integrated assembly/genetic map that are not fully resolved – and only in these regions – gene location data are added to the graph. For each set of unoriented scaffolds that belong to a particular node in the map, Chromonomer will find a consistent set of orthologous genes, and order and orient the scaffolds to match the order in the external genome. At least one ortholog must be present to order a scaffold, and at least two orthologs to orient a scaffold.

For the Gulf pipefish, we employed the genome of its congener, the greater pipefish (*Sygnathus acus*), to provide conserved synteny information. Figure 3 shows the result on Gulf pipefish linkage group 14 (LG14). After the initial incorporation of the genetic map (Fig. 3A), a number of scaffolds are not entirely resolved, including a large cluster of scaffolds at the top end of the chromosome (LG14, 0cM), along with several additional scaffolds at interspersed cM nodes. Provided scaffolds and conserved gene synteny information (Fig. 3A, colored boxes), Chromonomer was able to order 16 scaffolds and could also orient 14 scaffolds (Fig. 3B, colored boxes). Figure 4 shows conserved synteny between *S. scovelli* and *S. acus*, before and after Chromonomer employed ortholog-based ordering. Naturally, the process has made *S. scovelli* LG14 look more like *S. acus*, which might not always be biologically correct. If the reference organism is sufficiently closely related, however, this method provides a rational hypothesis for a likely order and orientation beyond what was initially arbitrary. This rationale is supported in this case by the fact that many of the reoriented scaffolds display conserved gene order also across their length, and genome-wide there is a strong pattern of conserved synteny between the two pipefish (Fig. S1). Figure 4 also shows that there remain putative true rearrangements between *S. scovelli* and *S. acus*, as demonstrated by Gulf pipefish scaffold 16 in the region at ~14Mb on LG14 (~22Mb in *S. acus*). In this case, the scaffold has high support from the genetic map for its position, while the orthologous gene block appears in a different relative location in the *S. acus* genome.

**Figure 3.**
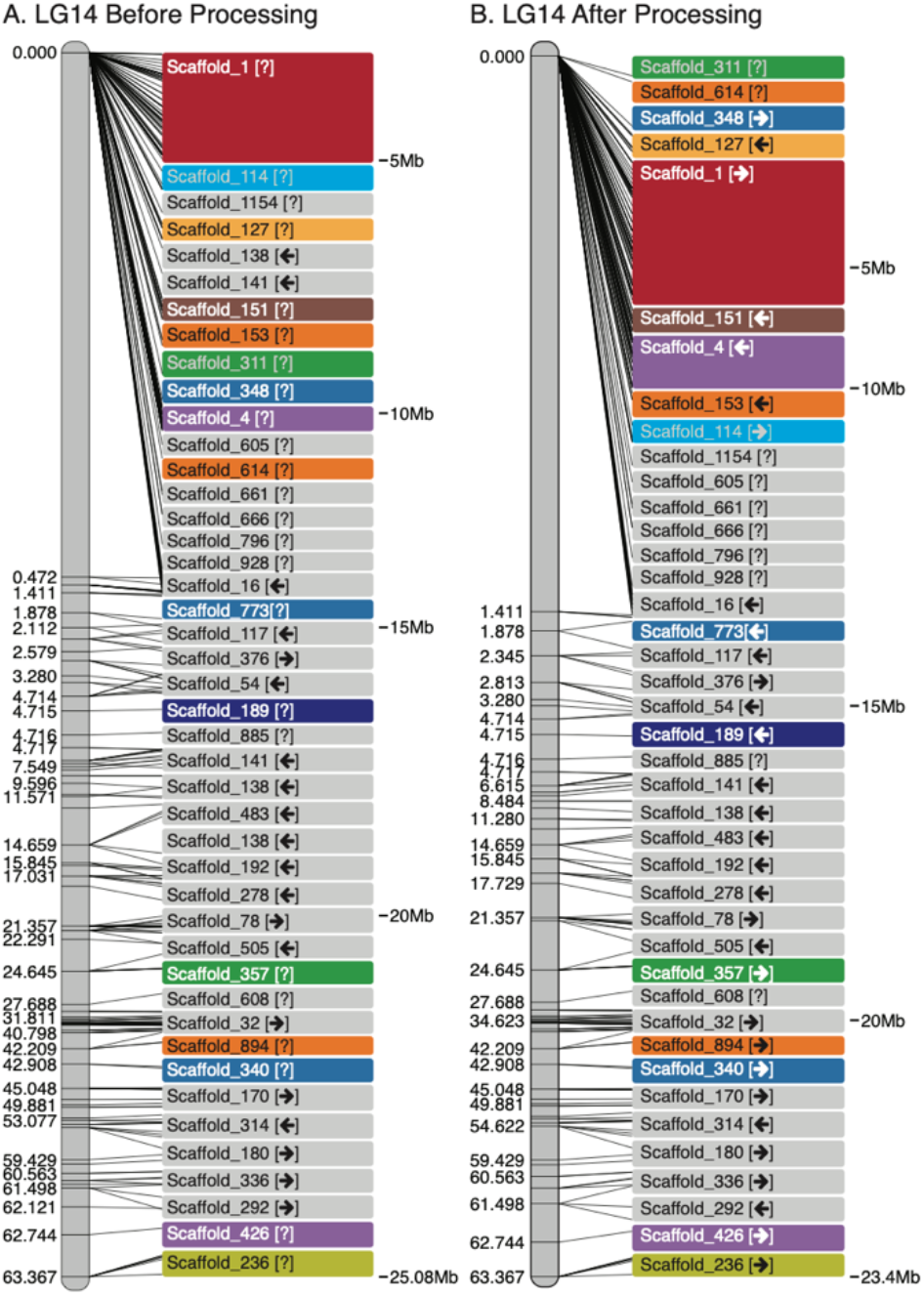
Including conserved gene synteny into the Chromonomer algorithm, as employed in the Gulf pipefish assembly. The figure shows LG14 before (A) and after (B) processing. In this example we have incorporated conserved gene synteny from the closely related *Sygnathus acus* to order and orient scaffolds whose position and orientation are left ambiguous by the genetic map. Colored scaffolds indicate where synteny was employed.

**Figure 4.**
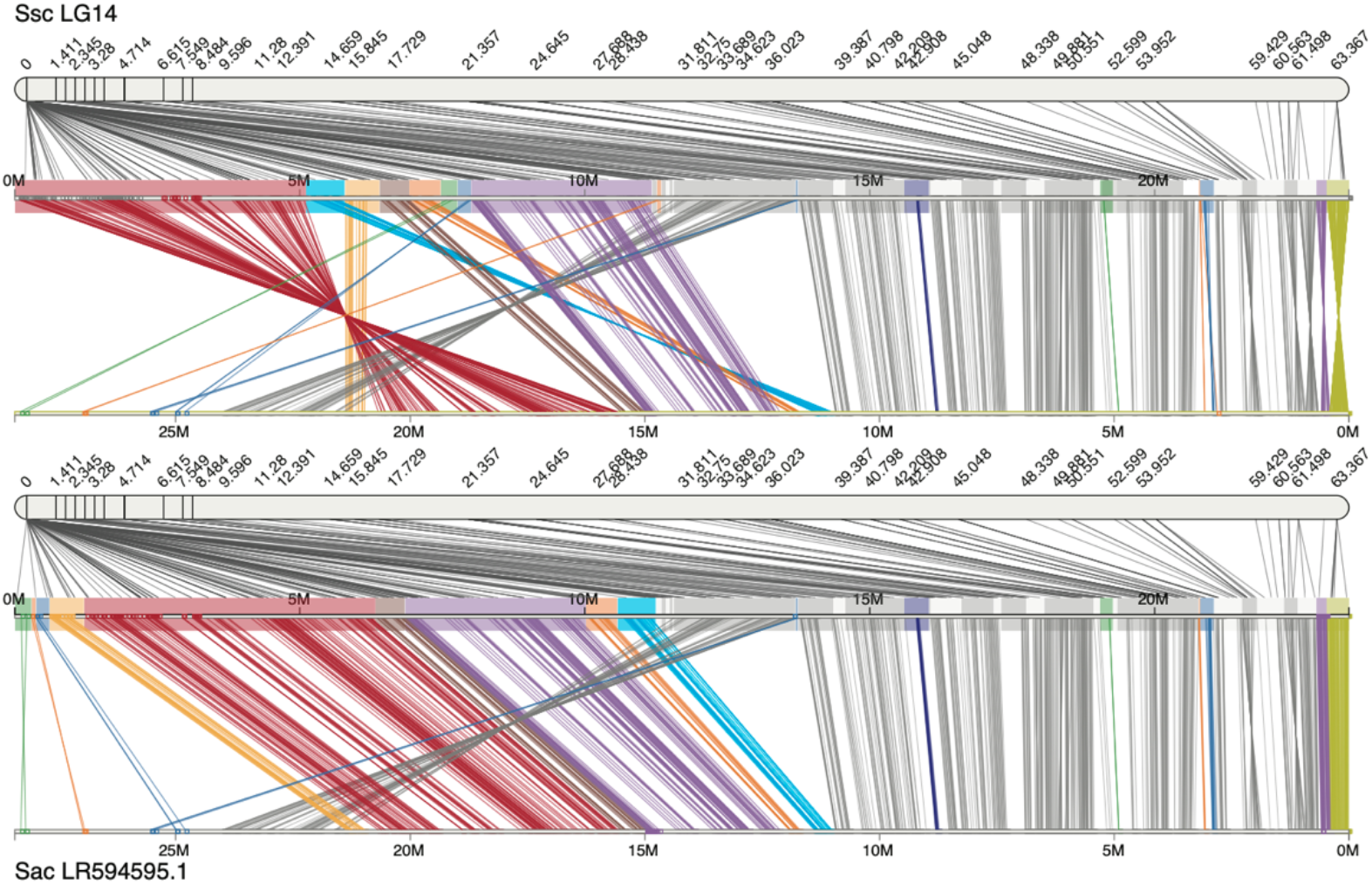
Ortholog-directed scaffold rearrangements in the Gulf pipefish. Potential improvements in LG14 integrated assembly by incorporation of gene synteny between *S. scovelli* and *S. acus*. Colored scaffolds indicate where synteny was employed, and colors are consistent with Figure 3. In each panel, the *S. scovelli* genetic map is shown above, linking the scaffolds of the physical assembly together. Lines also connect a pair of orthologous genes together between *S. scovelli* and *S. acus*.

### Identifying putative positions of misassembly using raw read coverage

Genome assemblers make errors during the assembly process, incorrectly joining and/or orienting scaffolds, often because assemblers are fooled by regions containing repetitive DNA sequence. With a hybrid or short-read-based assembly, scaffolds sparsely contain gaps between contiguous regions, inferred from mate-pair libraries or low-coverage long-reads. If the genetic map indicates a misassembly, Chromonomer relies on the largest gap between the defining markers to position where to break. However, the space between markers in conflict might encompass a large genomic region, or a repeat might be small or have enough identity to cause a mis-join without nearby gaps. If the assembly was constructed exclusively from long-reads, there might be no gaps present at all, despite the fact that hybridizing software may have silently reordered or reoriented regions of sequence, based on Hi-C or optical mapping data, for example. Yet independent evidence such as conserved gene synteny might insinuate that misassemblies are present. Additional methods are therefore needed to define breakpoints for adjusting the assembly. A reasonable proxy for the location of repeat regions in the genome are segments in the assembly with anomalous read depth, identifiable by aligning the raw reads back onto the assembly. Repeat regions can appear as segments of very high (or sometimes very low) coverage. If these depth of coverage data are provided to Chromonomer, it can use a sliding window algorithm on each scaffold to identify regions of high or low coverage – defined (by default) as three standard deviations above or below the mean per-scaffold coverage. Chromonomer will then use these peaks to insert virtual gaps into the integration graph, and these will become possible breakpoints when the Chromonomer integration algorithm executes (Fig. 5).

**Figure 5.**
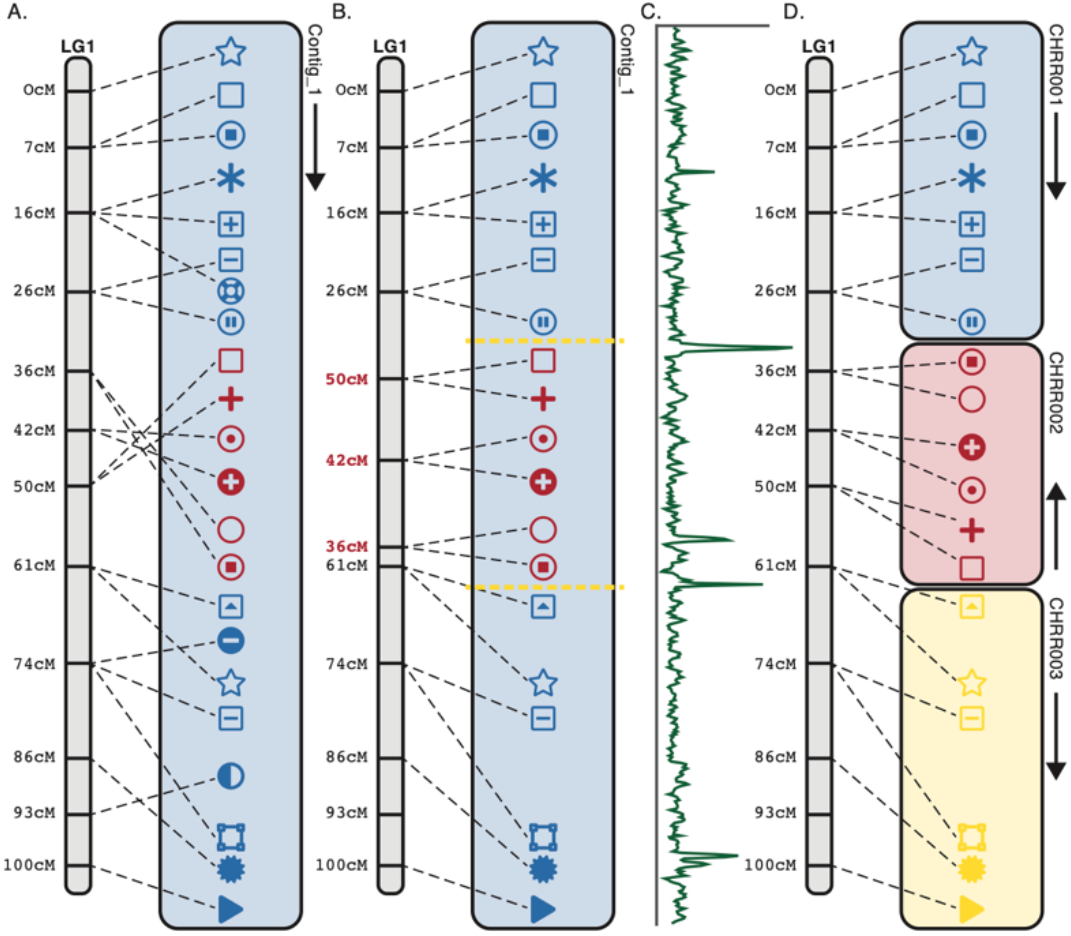
Using depth of coverage to create virtual gaps and rescaffolding the assembly. Assemblies constructed using long reads often consist purely of contigs, with no gaps. In these cases, we can input into Chromonomer depth of coverage data, generated by aligning raw reads back to the assembly, and we can identify anomalous values in depth of coverage to direct where to create virtual breakpoints in the assembly. Here, in (A) the markers clearly show a misassembly in the center region of the contig (red markers). With no gaps, the normal algorithm to split the contig will fail. (B) The scaffolding algorithm instead assumes the genetic map is the correct source of information and identifies where the contig should be broken, according to a consistent set of reordered map markers. Depth of coverage information (C) is incorporated to identify logical break points and (D) the contig is split into the respective pieces.

### Rescaffolding an assembly prior to map integration, as deployed in the platyfish genome

Chromonomer relies on scaffolds that are constructed from sets of contigs and gaps to properly integrate assemblies. In many long-read assemblies, such as produced by WTDBG2 (Ruan and Li 2019) there are no scaffold objects, only gapless contigs, while in others there are very few gaps (e.g., Flye (Kolmogorov et al. 2019)). These contigs tend to have robust assembly characteristics; nonetheless, misassemblies can still occur. As described above, the basal Chromonomer algorithm connects scaffolds (or contigs) to the linkage group graph and seeks a consistent order of markers between the linkage group and the set of scaffolds. However, if there is a misassembly that inverts or translocates a component of the contig but does not produce scaffold gaps, the basal algorithm on its own will discard all inconsistent markers but the majority set, leaving the scaffold located (and unbroken) at the place in the graph with the largest number of consistent markers. We can see how this would occur in the southern platyfish (*Xiphophorus maculatus*) assembly (Schartl et al. 2013). The assembly is qualitatively impressive, with 24 chromosome-length contigs, and only an additional 76 scaffolds, with an N50 of 31.5Mbp. The assembly was generated from 83x coverage from a PacBio Sequel I instrument, polished with Illumina short-read data, and an optical map (Bionano) was used to further scaffold the assembled contigs. However, when we compare the assembly against a high quality genetic map containing more than 22,000 markers and 267 progeny (a backcross between *X. maculatus* and *X. helleri* (Amores et al. 2014)), we find that while some putative assembled chromosomes agree strongly with the genetic map (linkage group 1, Fig. S2), others show potential misassemblies, such as linkage group 14 (LG14, Fig. 6A). (Since this genetic map was produced from a hybrid cross, it is a possibility that some of the conflicts between the assembly and map *could* be due to true differences between the *X. maculatus* and *X. helleri* genomes.) Here, the map shows a large inversion relative to the assembly on LG14 between ~35-53cM. As mentioned above, the basal Chromonomer algorithm prioritizes contig sequence over marker order, but it trusts marker order over scaffold sequence; that is, if a genetic map were to indicate an assembly error, and there exists a clear place to break the assembly, Chromonomer would break it, but if the sequence is contiguous, it assumes the marker is mis-genotyped and retains the original unbroken sequence. In platyfish LG14, Chromonomer would not be able to find a correlated maker order between the linkage group and the physical assembly so it would discard all conflicting markers until the largest set of correlated markers remained. Chromonomer would then place the should-be-split, unbroken contig at this final position.

**Figure 6.**
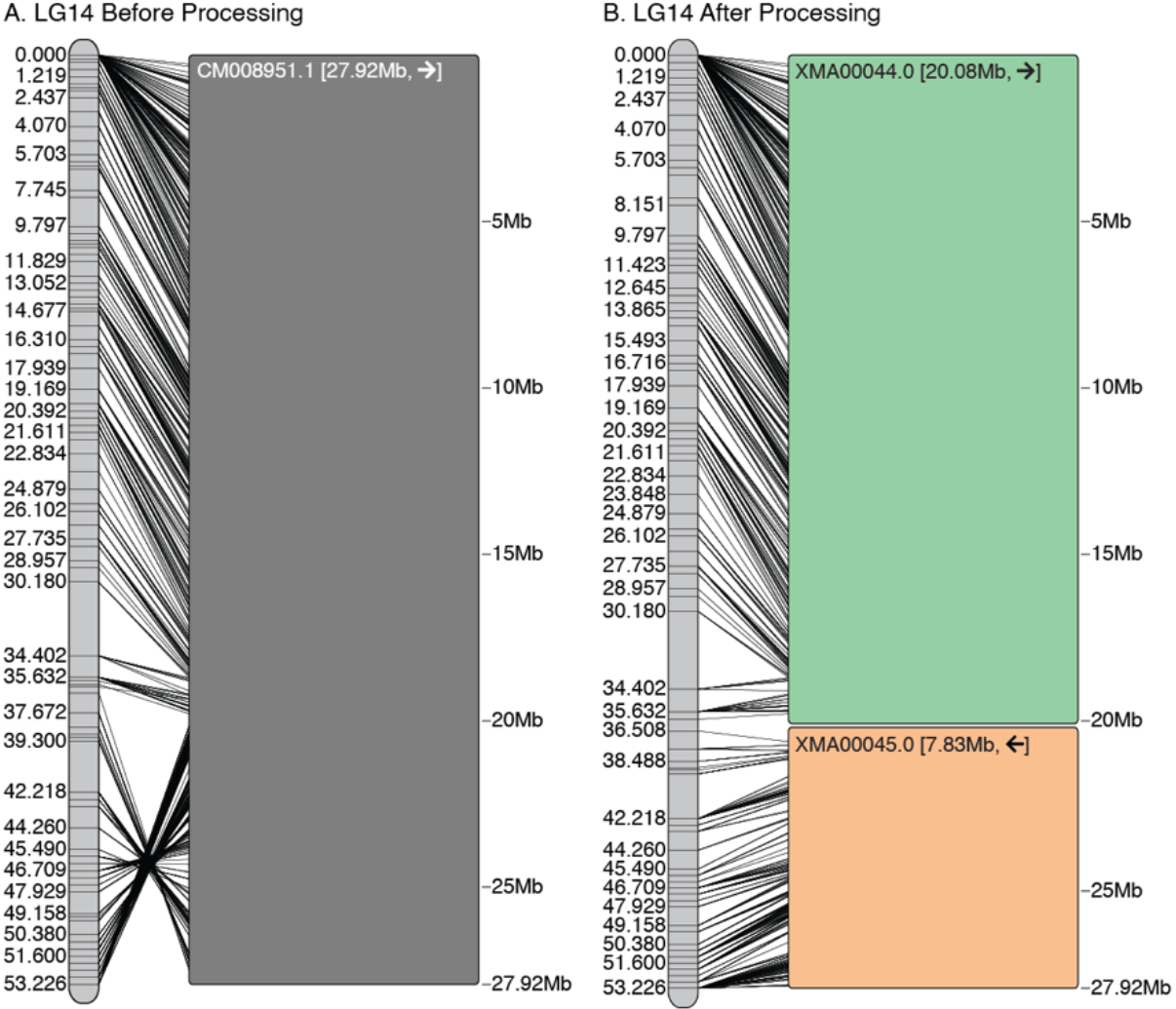
Using virtual gaps and the *rescaffold* algorithm in platyfish. A. The platyfish assembly shows a clear misassembly (inversion between ~35-53cM) when compared against the genetic map. B. A consistent order of markers is found on the map, and depth of coverage is employed to split the CM008951.1 contig into 2 components that can then be independently reoriented.

For such misassembled, gapless long-read contigs where assembly error is likely, Chromonomer provides the *rescaffold* option (Fig. 5). When the *rescaffold* algorithm is enabled, Chromonomer now prioritizes the genetic map order over the contiguous sequence assembly. In this model, Chromonomer assigns each map node (cM location) ownership of the physical sequence defined by its set of markers and that will allow map nodes to move ‘their’ associated sequence around in order to match the orientation and order specified by the genetic map. To do this, Chromonomer first must trim markers from the edges of each cM map node so that the physical positions of markers from any single node do not overlap with any other node (e.g., Fig. 5A, the physical position of two markers from the 16 and 26cM nodes overlap and must be rectified; see Methods for details). Once map nodes do not overlap in the physical sequence, Chromonomer will reorder the map nodes (Fig. 5B, red cM nodes) to match the order of markers in the physical assembly. Breakpoints are required for the algorithm to proceed further, if the sequence in the integration has no gaps, Chromonomer can insert virtual gaps via raw read depth of coverage (as described above, Fig. 5B, yellow lines). The physical assembly next is split, and the markers are returned to their original, cM map order (Fig. 5D); this results in the split contigs being ordered and oriented according to the genetic map. Figure 6B shows the result of applying this algorithm to LG14 of the platyfish assembly, where marker order is not cleanly correlated between the genetic map and physical assembly. We can examine the outcome by comparing patterns of gene synteny relative to the medaka genome (*Oryzias latipes*, Fig. 7) before and after running Chromonomer. The reordered physical assembly is congruent with gene order in medaka.

**Figure 7.**
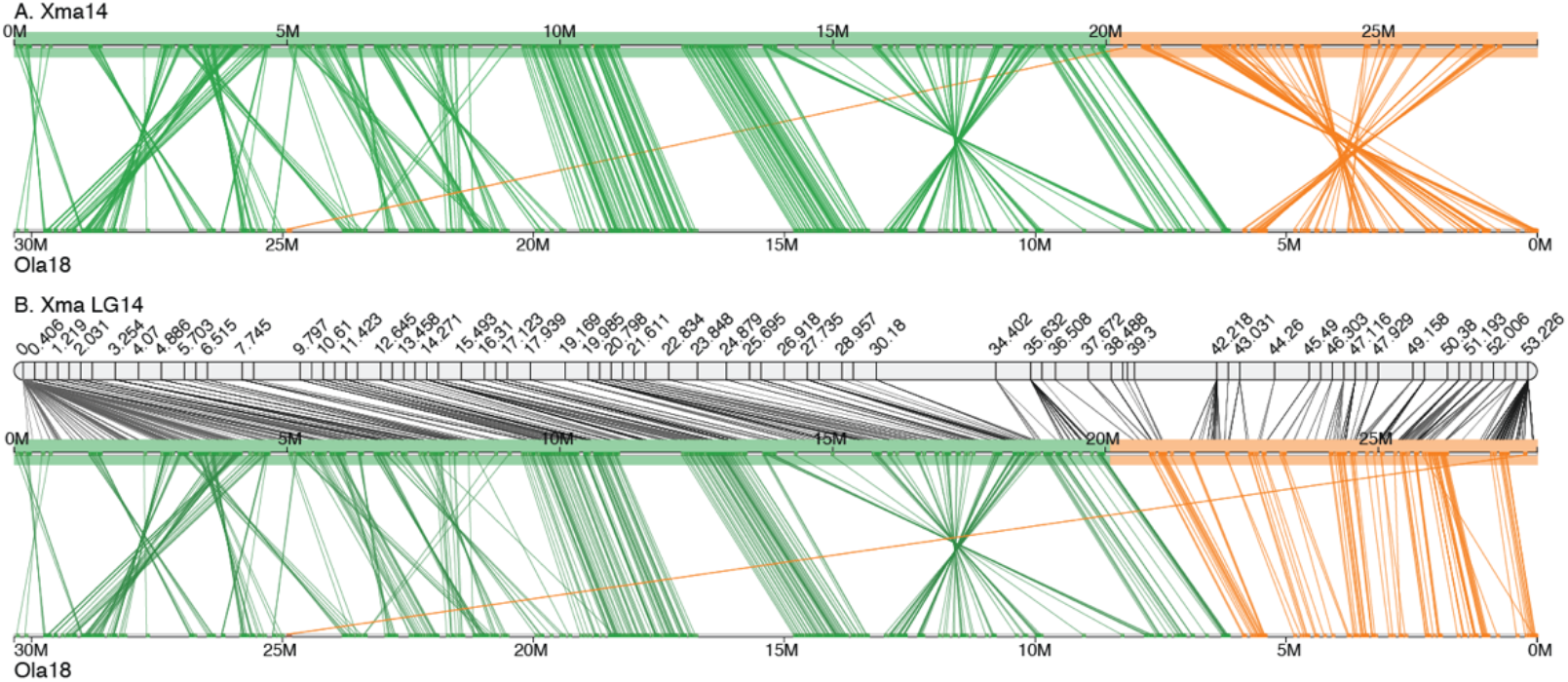
Improvements in the platyfish chromosome-level assembly. Conserved gene synteny between platyfish (Xma) and medaka (*Oryzias latipes*, Ola) illustrates improvements in the LG14 integrated assembly by application of the *rescaffold* algorithm. The top panel shows synteny prior to correction, several inversions are present, including one associated with the platyfish assembly (orange, colored to match the scaffolds in Fig. 6). After correction, inversions and ordering are rectified.

A more complex case is demonstrated on linkage group 3 (LG3, Fig. S3A). In this case, a chromosome-length contig is placed into the integration graph twice, with two subsets of markers attached to each node. A small scaffold (PGSD01000081.1) belongs in the middle of this chromosome, according to the genetic map. We also see at least two major inversions and several smaller errors in the assembly when we view the order of the markers in cM versus basepair coordinates. Chromonomer is able to apply the *rescaffold* algorithm and correct the ordering on LG3 (Fig. S3B) retaining 80% of the markers that were genetically and physically congruent in the final order. When we again examine the Chromonomer corrections using conserved synteny (this time with respect to the pufferfish, *Takifugu ruprides*, Fig. S4), the pattern is consistent with improvement in the integrated assembly.

### Algorithmic limits of Chromonomer demonstrated by the rockcod chromonome

The Antarctic bullhead notothen, or black rockcod (*Notothenia coriiceps*), is an extreme cold-adapted fish with an interesting karyotype. While the ancestral haploid chromosome number in teleost fish is 24 or 25 (Naruse 2004), the black rockcod has just 11 chromosomes (V. P. Prirodina, A. V. Neyelov 1984). Using an outcrossed RADseq-based genetic map constructed from 244 progeny in an F1 pseudo-testcross with 9,138 mappable markers, Amores et al. (2017) were able both to confirm this genome evolution occurred by end-to-end fusions and to identify which ancestral chromosomes became fused. The rockcod physical genome was also assembled using a hybrid strategy that mixed data from Illumina paired-end libraries, from 454-sequenced mate-pair libraries, and from limited PacBio RS II sequencing. In that assembly, the Illumina and 454 reads were integrated using the Celera assembler (designed originally for Sanger sequencing data; Myers (2000)), while the PacBio reads were incorporated using Gapfiller (Nadalin et al. 2012) and PBJelly (English et al. 2012). The resulting assembly was composed of 37,605 scaffolds, was 636Mbp in length, had a scaffold N50 of 218Kbp, and the largest scaffold was 28.8Mbp in length — a poor result that is not atypical for Illumina-based assemblies. We used Chromonomer to integrate the physical assembly with the genetic map for the first time. However, upon doing so, we found extreme discordance within the assembled scaffolds. For example, the second largest scaffold, KL668296.1 (27.5Mbp in length), contains 368 markers, but these markers were scattered in the genetic map across every one of the 11 linkage groups (Fig. S5). In fact, the four largest scaffolds map to all 11 linkage groups resulting in a remarkably disordered assembly. The pattern can be seen when gene orthologs are visualized in comparison with a related genome, in Figure 8. The x-axis shows genes from rockcod in grey (at bottom) and the corresponding orthologous genes in the blackfin icefish (*Chaenocephalus aceratus*) in red, located on the *C. aceratus* chromosomes (y-axis), but ordered according to the rockcod. Large scaffold KL668296.1 (boxed by a dashed line in Fig. 8A) spans nearly half of rockcod linkage group 1, but genes orthologous to those identified on this scaffold are found dispersed all over the *C. aceratus* genome, a condition unlikely to be biologically true. A multitude of other rockcod scaffolds are also probably chimeric, most likely due to assembly errors that stem from mate-pair libraries, with error amplified by the hybrid assembly and gap closing/scaffold extension algorithms that were optimized for maximal simple statistics (like N50), but not for accuracy. After running the basal algorithm of Chromonomer, scaffold KL668296.1 was broken down into 27 coherent pieces and those were reintegrated into their respective positions according to the genetic map. The resulting increased congruence in conserved gene synteny suggests structural improvement of the assembly (Fig. 8B), and importantly, the original signal of chromosome fusion in the rockcod is more cleanly resolved as a synteny is split between LG1 and LG4 in *C. aceratus*. If we view the genome-wide conserved gene synteny between rockcod and the blackfin icefish, we see similarly improved synteny, but still a lot of noise. Because of the finely granular nature of misassembly in this genome, there are not enough markers to fully correct all of the errors, and segments containing single or small numbers of genes remain incorrectly fused to other segments (e.g., Fig. S6); such cases likely account for many of the lines crossing to non-orthologous chromosomes in Fig. 9B.

**Figure 8.**
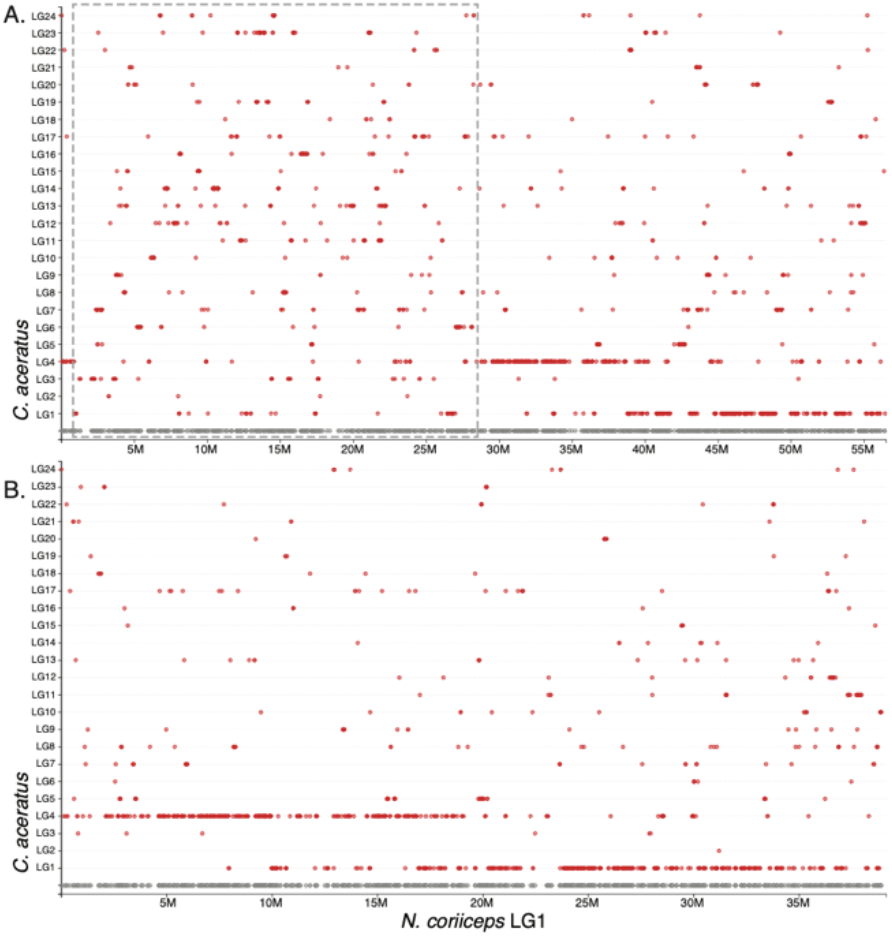
The *Notothenia coriiceps* assembly. All of the large scaffolds in the rocked assembly appear to be large chimeras from across the genome. When we examine LG1 in rockcod (A) we can see that orthologous rockcod genes are found across the related blackfin icefish assembly. The largest scaffold, KL668296.1 is highlighted by the dotted line and it is composed of sizeable pieces from all over the genome. (B) After processing with Chromonomer, the scaffold is broken up and redistributed in the assembly. We can now clearly see the conserved, two-to-one gene synteny between the icefish and rockcod.

**Figure 9.**
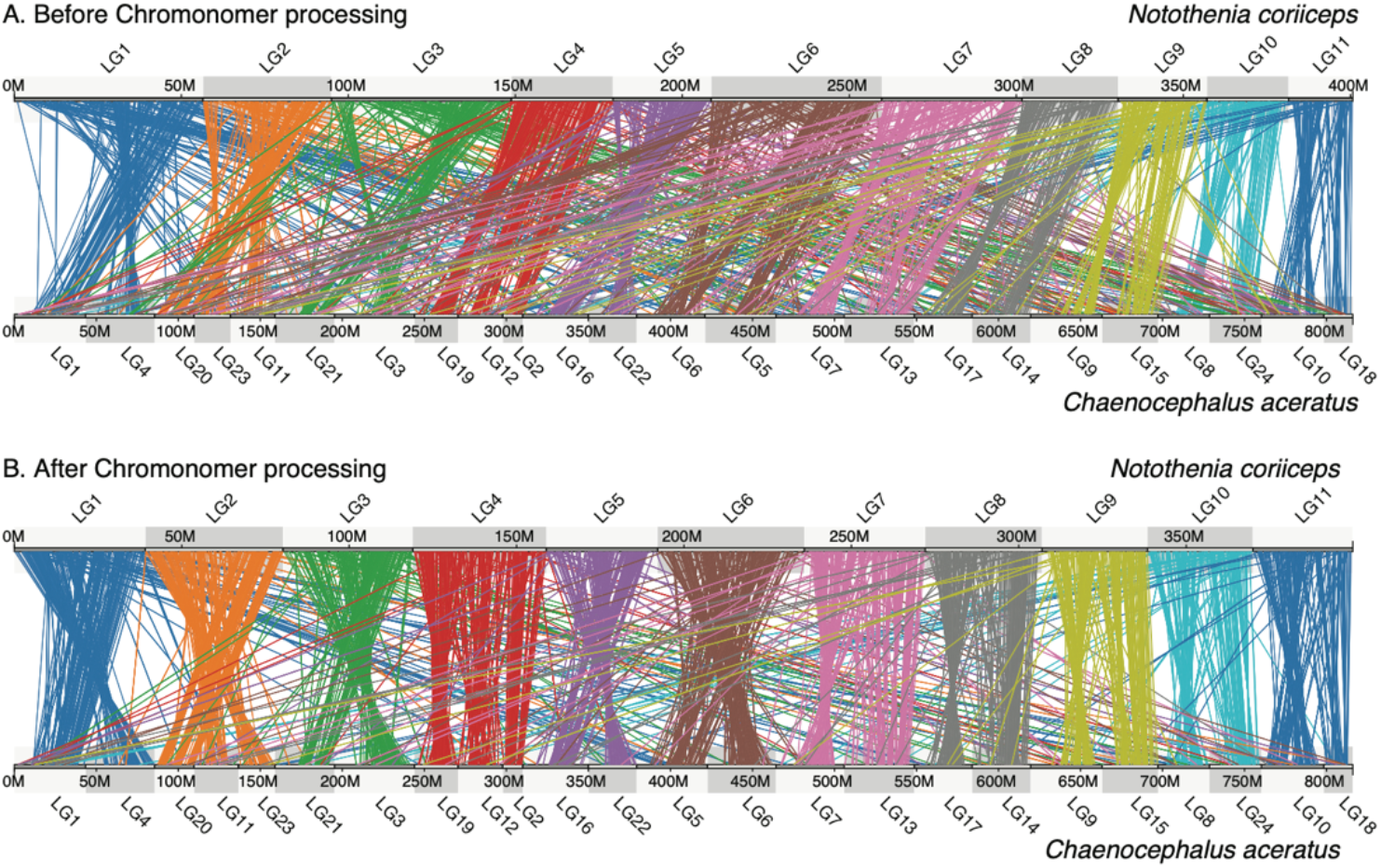
Chromonomer improves the rockcod assembly. The rockcod assembly can be chromonomed using the genetic map. (B) shows marked improvement in the assembly after breaking down the largest scaffolds using the genetic map. However, smaller assembly errors remain.

## Discussion

Physical genome assembly has only recently surpassed the pathbreaking human genome project (International Human Genome Sequencing Consortium 2001) in contiguous assembly length and precision. The human genome project was completed by a large consortium with massive resources (Collins 2003), but only a small number of other genome projects were conducted in an allied manner. With the arrival of short-read sequencing, genome projects have proliferated, resources for sequencing and assembly became diffusely distributed, and assembly quality commensurately dropped. Resources were rarely available to push a genome past draft form. Recently, sequencing technology has facilitated order of magnitude improvements to genome assembly though the employment of long-read, high volume sequencers. These technologies have finally surpassed the results achieved by a 1998 ABI Prism 3700 sequencer (Hutchison 2007), at vastly higher volume and lower cost. However, even with greatly improved N50s, accurately moving a genome from a collection of scaffolds to a chromonome (Braasch et al. 2015), with realistic long-range relationships among assembly segments, remains a major impediment.

Genetic maps are one of the oldest genomic resources, dating back to the beginning of the field of modern genetics (Painter 1933), but by the time of the human genome project they too were performed by only a small number of groups and required significant resources to discover and genotype markers. Short-read sequencing changed genetics too, as RAD sequencing and software like Stacks made genetic mapping broadly feasible and applicable (Rochette et al. 2019). This new generation of genetic maps could provide huge numbers of markers simultaneously with the genotyping itself, which allowed for new crossing designs to be completed in only a single generation. Genetic maps are generally created only for a subset of organisms that can be experimentally crossed, but F1 maps can be generated from crosses extracted directly from nature, and male-specific F1 maps can be produced even via single sperm sequencing (Xu et al. 2015; Sha et al. 2017)! Chromonomer can make use of any of these maps as well as those from standard, multigenerational crosses. Chromonomer expects a single map to be input to the program. To satisfy this requirement, maps can be joined together by the linkage mapper (e.g., the consensus map produced from F1 male- and female-specific maps), or a draft genome can be used to synthesize markers from different families together into a single map (Sutherland et al. 2016).

Chromonomer provides a method to integrate physical genome assemblies with genetic maps. Naively, this seems like a very simple process that should not require a dedicated piece of software. Each sequencing strategy, however, brings a particular error model along with it. A putative genome assembly is a hypothesis describing the actual, unknown genome. The key to successfully integrating a physical assembly with a genetic map is the ability to rank the quality of different types of information and to employ the most dependable information in the most rational hierarchy. Along these lines, Chromonomer is actually two distinct things: first, a tool to integrate physical and genetic assemblies, and second, a hypothesis generator to be employed during the assembly process itself.

To facilitate this process, Chromonomer provides the ability to integrate multiple lines of information. The basal algorithm compares the map positions of markers to their physical alignment positions with the goal of ordering and orienting scaffolds (or contigs) into chromosomes. The software commensurately provides extensive documentation, including web-based visualization drawings, to inform the user which genetic markers were discarded and why. Integrating the map and the physical assembly is meant to be an iterative process that allows the researcher to improve the map, a process in which genotyping errors become obvious in the context of the physical assembly. For example, if a postprocessing scaffold retains genes that, on the basis of conserved synteny, appear to belong elsewhere, the researcher could use the log that is generated from each scaffold integration to investigate whether Chromonomer removed a singleton marker that would otherwise have associated that segment with a location predicted by conserved synteny. In addition, poorly integrated genomic objects, such as from mate-pair sequencing, can create chimeras in a physical assembly, but Chromonomer can easily highlight these and correct at least a subset of them. The researcher, when presented with the number and type of scaffold splits conducted by Chromonomer, can also re-examine the various data types added to the assembly (e.g., a particular sequenced mate-pair library, or the output of software that hybridized different sequenced libraries together) and remove those that are generating large numbers of chimeras.

Another line of evidence that can be employed to improve assemblies is syntenic gene order conserved between closely related species. Chromonomer allows for this information to be employed when the genetic map is uninformative. On the one hand, using orthologous genes from another species to order scaffolds can make the focal genome artificially conform to that reference taxon. On the other hand, synteny is a powerful line of evidence and gene order is often conserved. In an iterative approach, a researcher can employ knowledge of synteny from multiple species to provide evidence for a particular assembly hypothesis. For example, if a change in gene order is contained by the boundaries of a scaffold, it is likely an assembly error (and vice-versa). As an example, we were able to enhance the assembly for Gulf pipefish by allowing Chromonomer to use gene orthology information from a congeneric species.

Incorporating raw read depth data into Chromonomer automates predicting where repeats such as transposable elements or satellites could have fooled the assembly algorithm and generated chimeras. Armed with this and Chromonomer’s *rescaffold* option, the researcher can further refine the elements of an assembly. Using the platyfish genome, we were able to correct multiple errors after the fact, even in a genome based on a long-read assembly. However, if we were actively performing the assembly *de novo* (instead of using it as an existing case study), we would be wise to go back to the optical mapping data and the assembler itself to investigate what initially caused the misassemblies that Chromonomer discovered.

Finally, the hypothesis-generating features of Chromonomer provide the researcher a means to test the success or failure of hybrid assembly strategies (such as the outcomes of automated gap-filling and scaffolding), as well as a means to test the robustness of the marker genotyping accuracy and marker placement in the genetic map itself. These features of Chromonomer mean the final integration of the genetic map and genome will be robust, as the researcher has been given the opportunity to correct underlying problems that would otherwise have been obscure.

In the decade where short-read sequencing dominated (such as when the rockcod reference genome was generated), the measures that were pursued to improve genome assemblies – employing hybrid assembly approaches alongside short-read sequencing – were largely unsuccessful. Many of the genomes in the literature today are of relatively low quality, and the assemblies could well be positively misleading. One measure of assembly quality is median N50 scaffold length, and its magnitude can be increased using hybrid assembly techniques while simultaneously creating low assembly accuracy (often due to mis-joins created by repeat regions that incorporate sequence multiple times, (Sohn and Nam 2016)).

Available assembly softwares still tend to operate as “black boxes”, with internal algorithmic decisions opaque to the outside user. Assemblers are still prized if they operate independently and akin to a home bread making machine – in go the raw ingredients and out pops a finished loaf. But if the bread doesn’t taste good, you probably would discard it instead of publishing your recipe. Further, many newer technologies to integrate long range information, such as Hi-C libraries (Lieberman-Aiden et al. 2009) or 10X Genomics libraries (Weisenfeld et al. 2017; Marks et al. 2019), include proprietary software and unpublished algorithms. Future work in accurate reference genome construction should focus first on exporting information that otherwise only the assembly software can know at the time it is executed, and second on identifying additional lines of evidence that can be properly integrated into an evolving assembly hypothesis.

## Methods

### The basal Chromonomer algorithm

Chromonomer requires a description of the genome assembly, which consists of an AGP file (NCBI 2019) describing the structure of the scaffolds (gap locations), a tab-separated file describing the genetic marker positions and source linkage group of each, and a file in SAM or BAM format (SAM/BAM Format Specification Working Group 2019) describing the alignment positions of the markers in the physical assembly. Optionally, a FASTA file containing the genome sequence can also be supplied (and with it Chromonomer will provide a reordered FASTA file of physical sequence after the Chromonomer algorithm completes). The contig and scaffold IDs must match among the input files.

In the first stage of the algorithm, Chromonomer resolves *inter-linkage group conflicts*. For each scaffold, it sorts aligned markers by linkage group (Fig. 1A-C), and if markers on a single scaffold belong to more than one linkage group, Chromonomer will attempt to split the scaffold. To do so, the markers must be in two or more consistently ordered sets, with an available gap (pre-existing or manufactured) between them. If such a configuration is not available, Chromonomer will discard the minority set of markers (Fig. 1C) until the scaffold can be split, or until a single, consistent set of markers remain. Split scaffolds are renamed (in a user-definable way) and the details of the process are logged.

For each scaffold, Chromonomer determines a provisional orientation by calculating a linear regression between the cM and basepair positions of the markers aligned to each scaffold. Although not all markers are yet consistently ordered, given a positive regression Chromonomer will orient the scaffold in the forward direction, or if a negative regression, the orientation will be in the reverse. This requires a scaffold to be present in at least two cM positions in the map.

Next, Chromonomer resolves *intra-linkage group conflicts*. A graph is constructed to represent each linkage group, with each cM position representing a node in the graph. Scaffolds are anchored to their respective nodes in the map; if a scaffold spans consecutive nodes, the nodes in the graph are collapsed together. If a scaffold occurs in more than one non-consecutive node, the scaffold is placed at each of these nodes (Fig. 1D). On each occurrence of a scaffold, Chromonomer identifies a maximal set of markers for the associated node that have a consistent order with respect to each other (marker base pair positions increase with map cM position, or the orders are inverted if in reverse orientation) (Fig. 1E). The markers that conflict with respect to each node are logged and discarded.

Now, the graph is destroyed and rebuilt from only the marker-pruned data using the same algorithm. Chromonomer traverses each graph and removes scaffolds that are not supported by *sufficient* markers (user definable, default is 1 marker). The software now looks at each scaffold individually within the linkage group. Importantly, if a scaffold was split because of an *inter-linkage group* conflict, it will be represented at this point in the algorithm as two or more separate scaffolds, which could potentially be split again to correct *intra-linkage group* conflicts (e.g., `CHRR0001` in Fig. 1B and 1D). Since the remaining markers must be consistently ordered, Chromonomer can now break scaffolds that are defined on more than one node in the linkage group at the nearest pre-existing or manufactured gap between the two groups of markers from each subset of the scaffold anchored to different graph nodes (e.g. `Scaffold_1` in Fig. 1E and 1F) and the details of each split are logged. If an appropriate gap cannot be found to split the scaffold, the smallest set of markers at a particular graph node are discarded until the scaffold can be split across graph nodes, or until there is only one set of consistent markers left in a single graph node, which places the unsplit scaffold in a single location.

Finally, Chromonomer recalculates the orientation of each scaffold, again using linear regression of marker positions, and summarizes the new, chromonome-level assembly, creating a new hybrid set of sequence (output in a FASTA file) and a new assembly description (output in a set of AGP files) to describe the changes. An external script, `translate_gtf.py`, is provided to lift over a set of gene annotations from a scaffold-based assembly to a chromonome, or vice versa.

### Depth of coverage and virtual gaps

Depending on the assembly process, a genome might be structured without any gaps. The lack of physical gaps does not imply that there are no misassemblies, however. A linkage map can reveal misassemblies (Fig. 6). Aligning raw reads back to an existing assembly can highlight likely areas of assembly ambiguity by revealing regions of anomalous depth of coverage. While there are many heuristics that can be applied to identify complex repeats, depth of coverage serves as a simple proxy for many of these. After aligning reads to a genome, samtools (Li et al. 2009) can be used to translate alignments to a per-base pair depth of coverage. These data can be specified to Chromonomer as a tab-separated file using the `--depth` option. Along each scaffold, Chromonomer slides a window (user definable, default 5Kbp) and calculates the mean depth of coverage within each window. It then determines how many standard deviations any window is from the scaffold mean. If a particular window is above or below the user-definable number of deviations (default is 3), a virtual gap of zero length is inserted at the 5’ end of the window into the internal AGP representation of the scaffold, which makes it available for Chromonomer’s standard splitting algorithm. Any virtual gaps not used during processing will be removed before outputting modified AGP or FASTA files.

### Ordering scaffolds with conserved gene synteny

If a scaffold does not span more than one cM node in the linkage group, it cannot be unarbitrarily oriented by the map. Likewise, if two or more scaffolds are collapsed into a single node, they cannot be unarbitrarily ordered. For these classes of scaffolds and only these, Chromonomer can be instructed to use the order of orthologous genes from a related genome to further specify order and orientation. In other words, conserved synteny data are subordinate to map location data. The user specifies the gene annotation of a related genome (in GFF or GTF format) using the `--orth_gtf` or `--orth_gff` flags, the annotation of the focal genome at a scaffold level using the `--gtf` or `--gff` flags, and the orthology assignment of genes between the two annotation files with the `--orthologs` flag (using a tab-separated file).

First, for each linkage group, the overall relative orientation of the external chromosome (i.e., from the related genome) must be determined. To do this, Chromonomer looks at all *anchored* scaffolds (existing on two or more nodes in the graph) in the focal genome and identifies the genes present. It then calculates the regression of gene positions between the linkage group and the orthologous genes on the external chromosome. The orientation of the external chromosome is reversed if it is determined by this method to be predominantly in the opposite orientation to the linkage group.

Next, for each node in the linkage group that contains more than one scaffold (that is a node in the map that hosts unordered or unoriented scaffolds), Chromonomer tabulates the genes present on the collection of scaffolds. Any genes that fall on a non-orthologous external chromosome or singleton, out-of-order genes are excluded, and the set of genes with a congruent order is retained. Similarly, orthologous genes that are too far away from the main set of orthologs, as determined using a trimmed mean algorithm (Bednar and Watt 1984), are discarded. Finally, the genes on the set of scaffolds are ordered according to the external chromosome position, which then allows ordering of the scaffolds in the node. Finally, the orientation of each scaffold is determined independently by calculating a regression based on the basepair positions of the genes on the scaffold and the external chromosome. Scaffolds that do not have any genes that are orthologous across both genomes, however, cannot be ordered or oriented by conserved synteny and they remained arbitrarily placed within the graph node (but after the ordered scaffolds).

### Rescaffolding based on the genetic map

For assemblies built from gapless, long-read contigs the basal Chromonomer algorithm could fail to correct misassemblies because incongruent marker orders have to be corrected by discarding markers (see basal algorithm above) within each contiguous sequence (since *contigs* are considered the most reliable information). However, the markers that would be discarded include the very markers that delineate the intra-contig misassembly. The *rescaffold* algorithm rescues situations like these by reprioritizing the map marker order over the basepair marker order. Each node in the linkage group graph will be considered the ‘owner’ of the sequence that ‘its’ markers span. The first step is to bucket sets of markers according to their graph node origin and to calculate a mean basepair position to represent the map node in the physical sequence. Next, the set of map nodes (and their markers) are re-sorted according to these mean basepair positions. Next, the map nodes are traversed in pairs, and the algorithm prunes any markers whose basepair positions overlap between the current and next map nodes. Markers are pruned in rounds, according to how far a marker is from the mean basepair position for the map node. This has the effect of removing markers that are the farthest away in the physical sequence from the mean map node position first. Pruning continues until no markers overlap between the nodes. Next, the contig is broken into pieces using the basal algorithms described above, and after splitting, the map nodes are re-sorted back to their cM positions; this constitutes the reordering and reorienting of the new components of the broken contig.

### Web-based visualization

Chromonomer creates ‘before’ and ‘after’ files for each linkage group, and for each scaffold that is modified, it creates a specific log for that scaffold. In addition, Chromonomer pre-computes a ‘before’ and ‘after’ JSON file (JavaScript Object Notation; https://tools.ietf.org/html/rfc8259) for each scaffold. General statistics, such as chromosome lengths, and the location and number of splits are logged. For each run of the software, this directory of data can be moved to a location visible to a web server and then served via HTTP. The JSON files are used to create visualizations of the ‘before’ and ‘after’ states of the linkage groups that can be viewed in a web browser (Figures 2, 3, and 6 are based on these visualizations). Chromonomer includes JavaScript code that will also be served by the web server to render these visualizations in the web browser using the D3 library (https://d3js.org/). Textual versions of these data are also available for local viewing if a web server is not available.

### Integration of the Gulf pipefish physical and genetic maps

The Gulf pipefish (*Sygnathus scovelli*) assembly (NCBI BioProject Accession PRJNA355893) is described in (Small et al. 2016) and was downloaded from https://creskolab.uoregon.edu/pipefish/. Briefly it was assembled from paired-end and mate-pair short-read Illumina libraries using ALLPATHS-LG (Gnerre et al. 2011). Small, et al. (2016) also describe the genetic map used in the assembly, with 108 progeny generated from an F1 cross using RADseq data assembled with Stacks (Catchen et al. 2011). An early version of Chromonomer was used to integrate the physical and genetic maps in Small, et al. (2016). For the present study, we aligned the RAD markers against the ALLPATHS-LG assembled set of scaffolds using BWA *mem* (Li 2013). To create the initial integration, we fed the tab-separated file containing the genetic map, the aligned markers in BAM format, the ALLPATH-LG FASTA sequences and AGP file describing the scaffold structure into Chromonomer, with default options. We then used the `translate_gtf.py` script, a Chromonomer accessory, to lift the gene annotations from the scaffold-only assembly to the chromonomed assembly.

### The addition of conserved gene synteny to the Gulf pipefish chromonome

To demonstrate the functionality of conserved synteny analysis, we used the newly available *Sygnathus acus* reference genome (unpublished, NCBI accession GCA_901709675.1, BioProject PRJEB32741), which is a highly contiguous, chromosome-level assembly generated from a synthesis of PacBio, 10X Genomics, and Dovetail Hi-C data. A gene annotation was not available, however. We took the raw RNAseq data from Gulf pipefish (Small et al. 2016) and aligned it against the *S. acus* assembly using STAR (Dobin et al. 2013). We then ran the BRAKER2 pipeline (Hoff et al. 2016, 2019) to create an annotation from the *S. acus* assembly, using the *S. scovelli* protein gene models, and the aligned *S. scovelli* RNA data. We then used the Synolog software (Catchen et al. 2009) to generate orthologs between *S. acus* and *S. scovelli*. Finally, we ran Chromonomer, now also supplying a scaffold-level gene annotation (in GFF format) for *S. scovelli*, the chromosome-level gene annotation from *S. acus* (generated by BRAKER2 in GTF format), and a tab-separated file listing the orthologs between the two genomes.

### Integration of the platyfish physical and genetic maps

The platyfish (*Xiphophorus maculatus*) assembly (NCBI accession GCA_002775205.2, BioProject PRJNA72525) is described in (Schartl et al. 2013). Briefly, it had been assembled by (Schartl et al. 2013) from PacBio Sequel data using HGAP4 (Chin et al. 2013), corrected with Arrow, and polished with Illumina data using Pilon (Walker et al. 2014). The genome was further scaffolded using a Bionano optical map resulting in a chromosome-level assembly. We employed the genetic map from (Amores et al. 2014) which was generated from an *X. maculatus* by *X. helleri* backcross resulting in 266 progeny, with markers generated by RADseq and assembled with Stacks (Catchen et al. 2011). We used a custom script to translate the scaffold names listed in the gene annotation GFF file to make them match those names in the assembly FASTA sequence and AGP files. We aligned the RADseq markers against the *X. maculatus* scaffold-level reference genome using BWA *mem* (Li 2013) and ran the basal Chromonomer algorithm, feeding the AGP, FASTA, and BAM files along with a tab-separated file containing the genetic map marker positions.

### Platyfish virtual breakpoints and the *rescaffolding* integration

We used the raw PacBio reads the *X. maculatus* assembly was based on from the NCBI Sequence Read Archive (accessions SRR7207855 – SRR7207868) and aligned them against the reference genome using Minimap2 (Li 2018). We then converted the alignments into measures of depth of coverage using the samtools *depth* module (Li et al. 2009). We ran Chromonomer again, supplying the depth of coverage as a compressed tab-separated file, and set the sliding window size to 1000bp and the depth of coverage standard deviation to 1 while enabling the *rescaffold* option.

### Integration of the rockcod physical and genetic maps

The black rockcod (*Notothenia coriiceps*) assembly (NCBI accession GCA_000735185.1, BioProject PRJNA66471) was created by (Shin et al. 2014) from PacBio RSII, Illumina HiSeq, and 454 GLX II data using the Celera assembler (Myers 2000), and gaps were filled and extended using the PacBio data with PBJelly (English et al. 2012). We employed the genetic map from (Amores et al. 2017) of 244 F1 progeny, with markers generated by RADseq and assembled with Stacks (Catchen et al. 2011). We aligned the markers against the black rockcod genome using BWA *mem* (Li 2013), used a custom script to translate scaffold and contig names in the NCBI GFF file to match the names in the NCBI FASTA and AGP genome description files. We ran Chromonomer, supplying the aligned markers in BAM format, the assembly AGP file describing scaffolds and contigs, and the FASTA sequence file, along with the marker positions in the genetic map in a tab-separated format.

## Data Access

All input data were previously available in online repositories and the appropriate accession numbers are listed in the Methods section.

## Acknowledgments

We thank Clay Small and John Postlethwait for comments on the manuscript. J. Catchen was supported by NSF grant 1645087. A. Amores was supported by NIH grant R01 ODO11116 (Postlethwait, PI) and NSF grant 1543383 (Postlethwait, PI). S. Bassham was supported by NIH grant R24 RR032670 (Cresko, PI). J. Catchen and A. Amores designed Chromonomer. J. Catchen implemented the software and completed the analyses. J. Catchen, S. Bassham, and A. Amores interpreted the analyses. J. Catchen and S. Bassham wrote the manuscript.

